# Metformin lowers HMGA1a and RBM3 protein in breast cancer cells, and its effect on *ESR1* splicing depends on cell density and exposure time

**DOI:** 10.64898/2026.11.04.764423

**Authors:** Yoshihiro Harada, Tomoko Tanaka, Yuriko Hamaguchi, Yuta Horita, Hiroyoshi Harada, Masayoshi Mori, Yusuke Murata, Kenji Ohe

**Affiliations:** Department of Clinical Pharmacotherapy, Faculty of Pharmaceutical Sciences, Fukuoka University, 8-19-1 Nanakuma, Jonan-ku, Fukuoka 814-0180, Japan; Department of Endocrinology and Diabetes Mellitus, Faculty of Medicine, Fukuoka University, 7-45-1 Nanakuma, Jonan-ku, Fukuoka 814-0180, Japan

**Keywords:** metformin, estrogen receptor alpha, ERα46, alternative splicing, tamoxifen resistance, HMGA1a, RBM3, cell density, breast cancer

## Abstract

The *ESR1* gene produces ERα46, an N-terminally truncated isoform generated by skipping of exon 1, which is reduced in tamoxifen-resistant breast cancer cells. We have reported that metformin moves the ratio of full-length ERα to ERα46 in opposite directions in MCF-7 cells and their tamoxifen-resistant subline TAMR1, and that metformin reduces TAMR1 xenograft growth (Harada et al., Am J Pharm Health Res 2022;10(11)). That report leaves two questions open: whether the effect is dose-dependent and stable across culture conditions, and what splicing factor might carry it. Here we report the experiments that bear on both.

Across a full dose range from 0.25 to 5 mM at high seeding density, metformin produced no consistent change in the full-length share of *ESR1* transcript in either line: densitometry of the gel gave 42%, 43%, 42%, 43%, 45% and 48% in MCF-7 cells and 55%, 44%, 52%, 47%, 68% and 43% in TAMR1 cells. The reported shift therefore depends on the seeding density used, and the comparison that produced it is not reproduced at high density. At low density the MCF-7 response persisted to 48 h (52%, 62% and 68% at 0, 2.5 and 5 mM).

Metformin lowered HMGA1a protein in MCF-7 cells and showed the same tendency in TAMR1 cells, at both seeding densities, and lowered RBM3 protein in TAMR1 cells at 24 and 48 h. HMGA1a is the factor we previously showed to generate ERα46 by trapping U1 snRNP at an upstream pseudo-5′ splice site, so a fall in HMGA1a predicts less ERα46, the opposite of what was seen in TAMR1 cells at low density. The fall in RBM3, the protein through which metformin alters splicing in myotonic dystrophy, is the more plausible candidate.

RT-PCR of the xenograft tumours from the earlier study detected full-length *ESR1* transcript in the untreated and tamoxifen-plus-metformin groups but not in the single-agent groups, with amplification failing in several lanes; these panels do not separate the treatments.

Taken together, the direction of the metformin effect on *ESR1* splicing is conditional on culture density and does not survive a dose-response test at high density, and HMGA1a is unlikely to be the factor that carries it.

## Introduction

About 70% of estrogen receptor (ER)-positive breast cancers respond to anti-estrogen therapy [1], but roughly one third relapse within 15 years [2]. Activating mutations in the ligand-binding domain of *ESR1* were identified in hormone-resistant metastatic disease [3,4], yet these account for only a fraction of tamoxifen-resistant tumours, so other mechanisms must operate.

One candidate is alternative splicing of *ESR1* itself. The gene produces ERα46, an isoform transcribed from the upstream non-coding exon E/F that skips exon 1 and therefore lacks the N-terminal AF-1 domain [5,6]. ERα46 antagonises the ligand-independent AF-1 activity of full-length ERα66, and its expression is reduced in tamoxifen-resistant breast cancer cells; re-expressing it inhibits proliferation [7]. Because ERα46 retains the C-terminal domain recognised by the antibodies used for clinical ER scoring, a tumour shifted toward ERα46 is still called ER-positive while behaving differently. We have shown that the splicing factor HMGA1a generates ERα46 by trapping U1 snRNP at an upstream pseudo-5′ splice site [8,9].

Metformin, a biguanide used widely in type 2 diabetes, modulates alternative splicing in vitro and in vivo, acting in part through a reduction in the RNA-binding protein RBM3 [10]. We recently reported that metformin at 2.5 and 5 mM moves the full-length/ERα46 ratio in opposite directions in MCF-7 cells and in their tamoxifen-resistant subline TAMR1, that the effect on ERα protein is tamoxifen-independent, and that metformin reduces tumour weight in TAMR1 but not MCF-7 xenografts [11].

That report establishes the phenomenon at two concentrations and one culture condition, and it does not identify a splicing factor through which metformin might act. Two questions follow directly. First, does the effect hold across a full dose range and across culture conditions, or is it specific to the condition tested? Second, does metformin change the abundance of HMGA1a, the factor known to produce ERα46, or of RBM3, the factor through which metformin alters splicing elsewhere?

Here we report a metformin dose-response from 0.25 to 5 mM at high seeding density, the 48 h time point at low density, immunoblots for HMGA1a and RBM3 in both lines, and RT-PCR of the xenograft tumours from the earlier study. None of these data appear in [11].

## Materials and methods

### Reagents and antibodies

Tamoxifen (Cayman Chemical, 13258) was dissolved in ethanol. Metformin hydrochloride (Tokyo Chemical Industry, M2009) was dissolved in water. Primary antibodies were anti-ERα (HC-20, sc-543; Santa Cruz Biotechnology), anti-HMGA1a (FL-95, sc-8982; Santa Cruz Biotechnology), anti-RBM3 (ab134946; Abcam) and anti-ACTB (A00702; GenScript). Blots were developed with SignalFire ECL reagent (Cell Signaling Technology, 6883S).

### Cell culture

MCF-7 breast cancer cells (ATCC HTB-22) were cultured in DMEM (Nacalai Tesque, 08456-65) with 10% EquaFETAL (Atlas Biologicals, EF-0500-A), non-essential amino acids and penicillin-streptomycin. The tamoxifen-resistant derivative MCF-7 Tam1 (ATCC CRL-3435, referred to here as TAMR1) was cultured in DMEM/F-12 (Sigma-Aldrich, D6434) with 1% EquaFETAL, 2.5 mM L-glutamine, 6 ng/mL insulin and penicillin-streptomycin. TAMR1 cells were maintained in 1 µM tamoxifen, which was removed where the experiment does not indicate it. Cells were grown at 37 °C in 5% CO_2_ and were used at log phase. Cell culture and compound conditions follow those of our earlier report [11].

Because the direction of the splicing response proved to depend on how densely the cells were plated, seeding density is stated for every experiment: cells were plated at either 1 × 10^4^ cells per well (low density, about 50% confluent at harvest) or 1 × 10^5^ cells per well (high density).

### RT-PCR

Total RNA was extracted from PBS-washed cells disrupted in TRIzol (Invitrogen, 15596026), precipitated from the aqueous phase and digested with RQ1 RNase-free DNase (Promega, M6101). One microgram was reverse transcribed with ReverTra Ace (Toyobo, 05038-64) and random primer (Toyobo, FSK-301), and PCR was performed with Ex Taq (Takara Bio, RR001A). Primers are those of our earlier reports [8,9,11] and are listed in Table 1.

**Table 1.**
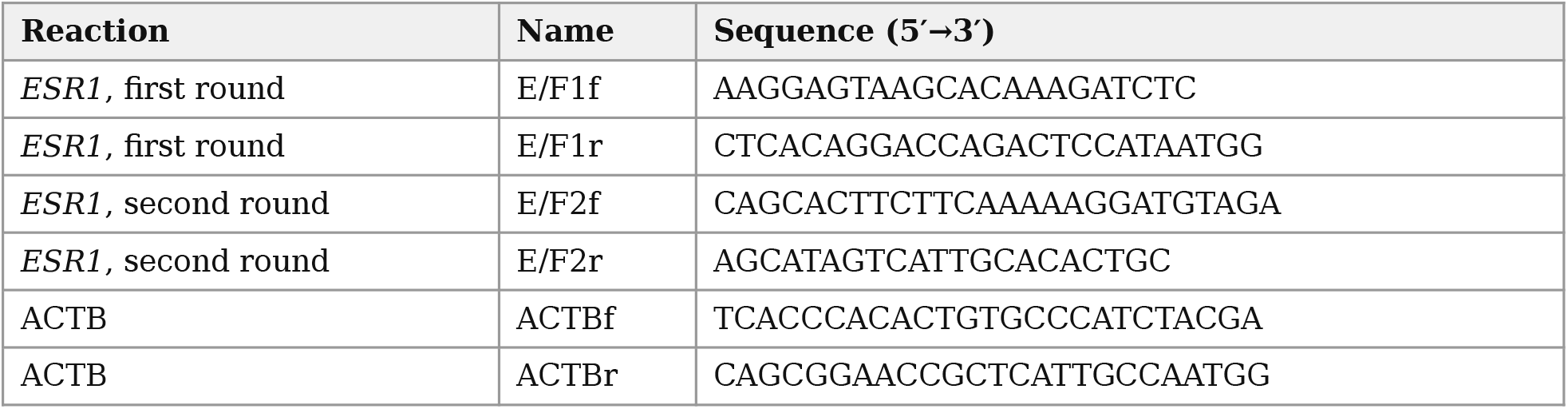
Primers used for RT-PCR. The two *ESR1* primer pairs are used in one nested reaction that ampli?es both the full-length and the ERa46 product; they are not pair-speci?c for one isoform.

The two *ESR1* transcripts were amplified together by nested PCR, as described for the same primers [9]. The first round used E/F1f and E/F1r for 20 cycles and the second round E/F2f and E/F2r for 25 cycles. Both rounds were run at 98 °C for 10 s, 60 °C for 30 s and 72 °C for 30 s. The nested reaction gives a full-length ERα product of 702 bp and an ERα46 product of 180 bp [9]; the difference is exon 1, which is retained in the full-length transcript and skipped in ERα46. ACTB was amplified in a separate single-round reaction under the same cycling profile, giving a product of 295 bp. Products were resolved on agarose gels alongside a DNA size marker.

Band intensities were quantified in ImageJ and the proportion of full-length transcript was calculated as full-length / (full-length + ERα46). We report this ratio as the full-length share throughout; the earlier report [11] gives the complementary ERα46 share, so the two sets of numbers sum to 100%.

### Western blotting

Protein was recovered by acetone precipitation from the interphase remaining after TRIzol/chloroform extraction, dissolved in urea-containing sample buffer (10 mM Tris-HCl pH 7.5, 25 mM KCl, 5 mM MgCl_2_, 6 M urea) and sonicated to shear co-precipitated nucleic acid. Protein was quantified with Pierce 660 nm Protein Assay (Thermo Fisher Scientific). Approximately 20 µg per lane was separated by SDS-PAGE and transferred to PVDF membrane by tank blotting. Membranes were blocked with Blocking One (Nacalai Tesque), incubated with primary antibody (1:1000) at 4 °C for 16 h and secondary antibody (1:5000) at room temperature for 1 h, and developed with Chemi-Lumi One L (Nacalai Tesque). Band intensities were quantified in ImageJ.

### Xenograft experiment

The tumours analysed here are those of our earlier report [11], and the animal work is described there. In brief, female BALB/c nude mice (4–8 weeks, Charles River Laboratories Japan) were implanted with 5 × 10^6^ MCF-7 or TAMR1 cells in Matrigel (Falcon, 354248). Each mouse received a tumour on each flank, in one of three combinations — MCF-7 on both sides, MCF-7 on one side and TAMR1 on the other, or TAMR1 on both sides — and the mice were identified by ear mark. Metformin (Wako Pure Chemical Industries) was given in the diet at 750 mg/kg/day and tamoxifen at 20 mg/day by gastric gavage, for four weeks. At the end of treatment a portion of each tumour was taken for RNA extraction and analysed by RT-PCR as above. All animal protocols followed National Institutes of Health guidelines and were approved by the Fukuoka University Committee for Animal Care (Protocol No. 171116, accepted April 2019).

### Reanalysis of public confluency RNA-seq

Scaled gene counts for GSE249290 [16], in which HEK293, A375 and SH-SY5Y cells were sequenced at about 50% and at 100% confluence in triplicate, were downloaded from the Gene Expression Omnibus. The deposited matrix is labelled as scaled but is not centred between samples: taken as supplied, 87.3% of tested genes in HEK293 and 97.0% in SH-SY5Y have a negative log_2_ fold change at high density, which no biological account explains. Size factors were therefore recomputed by the median-of-ratios method on genes with a mean count of at least 10, after which the proportion of negative changes was 49.3%, 49.5% and 50.8% in the three lines and the median log_2_ fold change was within 0.011 of zero. Groups were compared by Welch’s t test on log_2_ normalised counts, p values adjusted by Benjamini-Hochberg across all tested genes in each line, and the three lines combined by Stouffer’s method on signed z scores. One high-density SH-SY5Y replicate had a size factor of 0.143; the analysis was repeated without it. Code and the full per-gene output accompany this report.

### Search for metformin datasets in oestrogen receptor-positive cells

The Gene Expression Omnibus was queried for all human metformin RNA-seq series (80 series, searched 5 November 2026), and the cell type and the processed files of each were inspected.

### Statistics

No statistical comparisons are made on the experimental data in this report. The band ratios are the values measured in the experiment shown, and the number of repeats behind each is stated in the figure legend. The tests named above apply only to the public-data reanalysis.

## Results

### A full metformin dose range produces no consistent change at high cell density

MCF-7 and TAMR1 cells were treated for 24 h with metformin from 0.25 to 5 mM at high seeding density and *ESR1* transcripts were amplified by RT-PCR (Fig. 1). Both products were present at every concentration. Densitometry of the gel gave a full-length share in MCF-7 cells of 42%, 43%, 42%, 43%, 45% and 48% at 0, 0.25, 0.5, 1, 2.5 and 5 mM: a rise of 6 percentage points at the highest concentration, with no change over the lower four. In TAMR1 cells the values were 55%, 44%, 52%, 47%, 68% and 43%, varying by 25 percentage points with no relation to dose.

**Figure 1.**
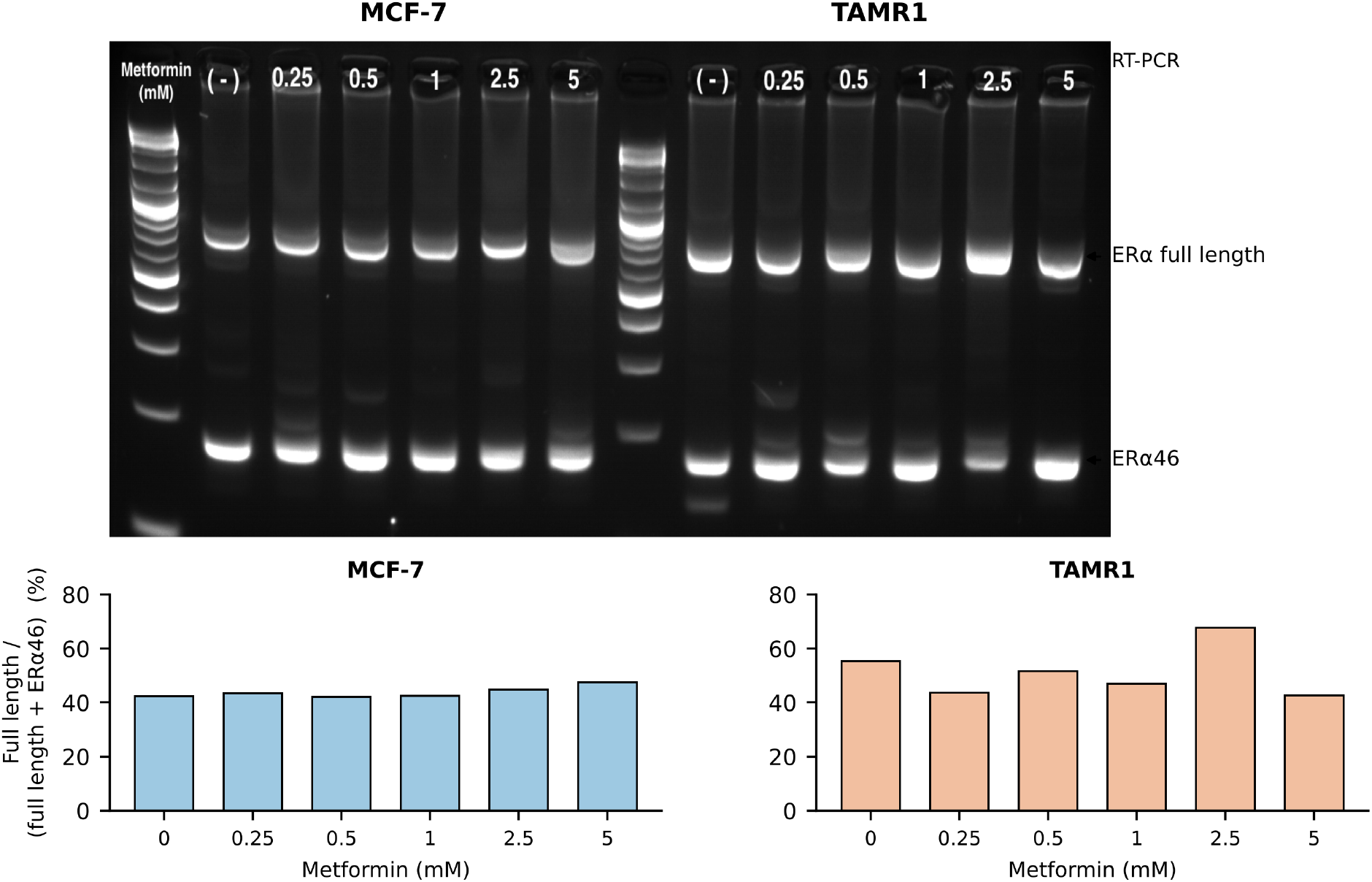
A full metformin dose range produces no consistent change in *ESR1* splicing at high cell density. RT-PCR of *ESR1* transcripts in MCF-7 (left) and TAMR1 (right) cells seeded at high density and treated for 24 h with 0, 0.25, 0.5, 1, 2.5 or 5 mM metformin, with a DNA size marker between the panels. Arrowheads mark the full-length ERα product and the ERα46 product. Below, the full-length share, full-length / (full-length + ERα46), obtained by densitometry of the gel shown with local background subtracted from each lane. Single experiment; no statistical comparison is made. These values were measured from the gel image for this report because the original quantification could not be located, and are provided as a source-data file.

At 2.5 and 5 mM, the concentrations at which we previously reported a shift [11], the high-density values here are 45% and 48% in MCF-7 cells and 68% and 43% in TAMR1 cells. The large opposite-direction shift reported at low density is therefore not reproduced at high density, and no dose-response underlies it.

### The MCF-7 response at low density persists to 48 h

The 24 h low-density experiment is published [11]. At 48 h under the same conditions (Fig. 2), the full-length share in MCF-7 cells was 52%, 62% and 68% at 0, 2.5 and 5 mM, the same direction as the published 24 h values of 58%, 57% and 74%. In the TAMR1 48 h gel only the untreated lane gave an interpretable pair of bands (56%); the 2.5 mM lane resolved into multiple products and the 5 mM lane did not amplify, so no 48 h comparison can be made for that line.

**Figure 2.**
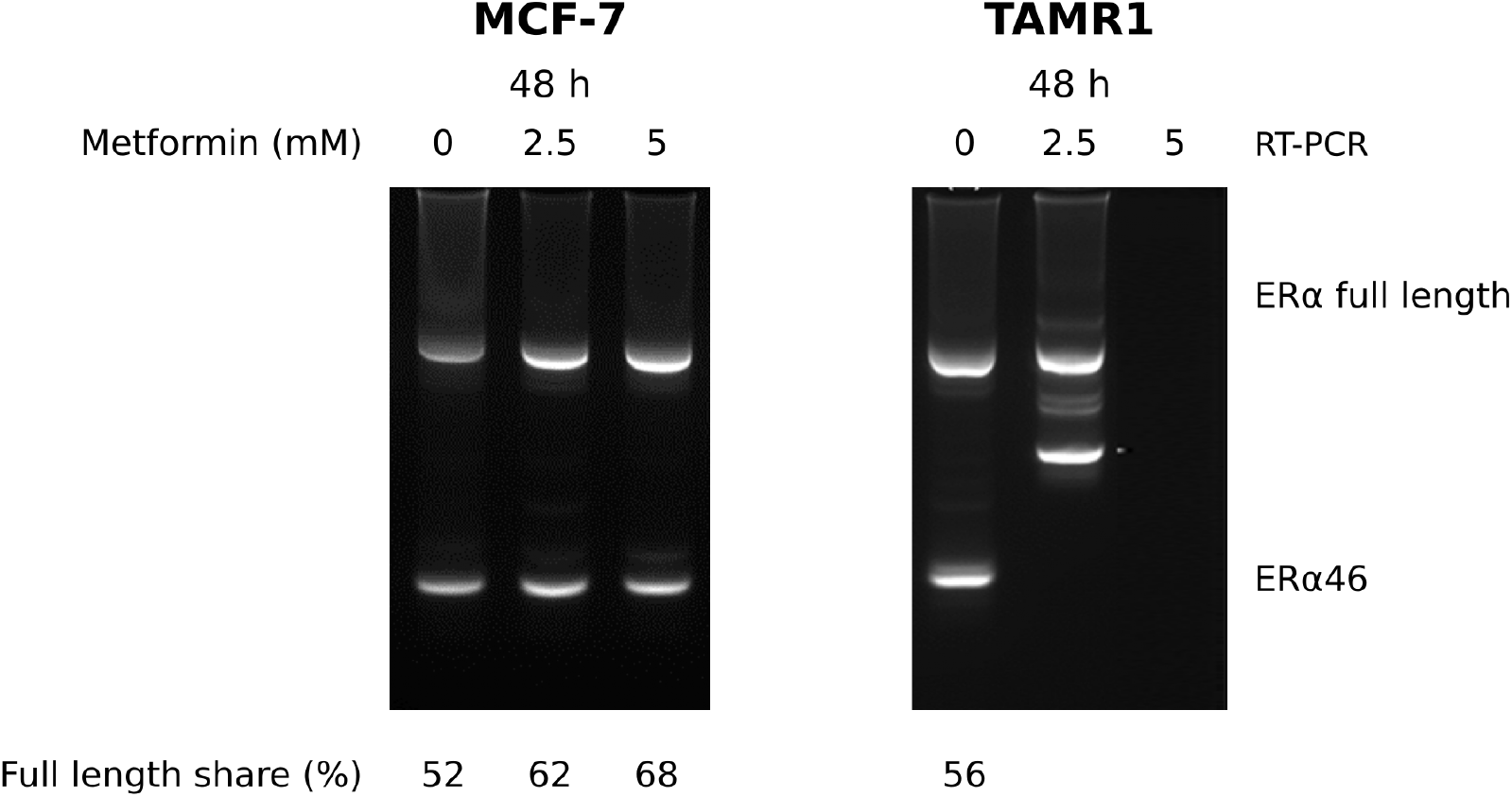
The MCF-7 response at low cell density persists to 48 h. RT-PCR of *ESR1* transcripts in MCF-7 (left) and TAMR1 (right) cells seeded at low density (about 50% confluent at harvest) and treated with 0, 2.5 or 5 mM metformin for 48 h. Numbers below the lanes give the full-length share as a percentage. In the TAMR1 panel only the untreated lane gave an interpretable pair of bands. The corresponding 24 h experiment is published [11] and is not reproduced here. Single experiment.

### Metformin reduces HMGA1a and RBM3 protein

HMGA1a protein decreased with metformin in MCF-7 cells and showed the same tendency in TAMR1 cells, at both seeding densities (Fig. 3). RBM3 protein, examined in TAMR1 cells at 24 and 48 h, also decreased with metformin. ACTB was used as a loading control.

**Figure 3.**
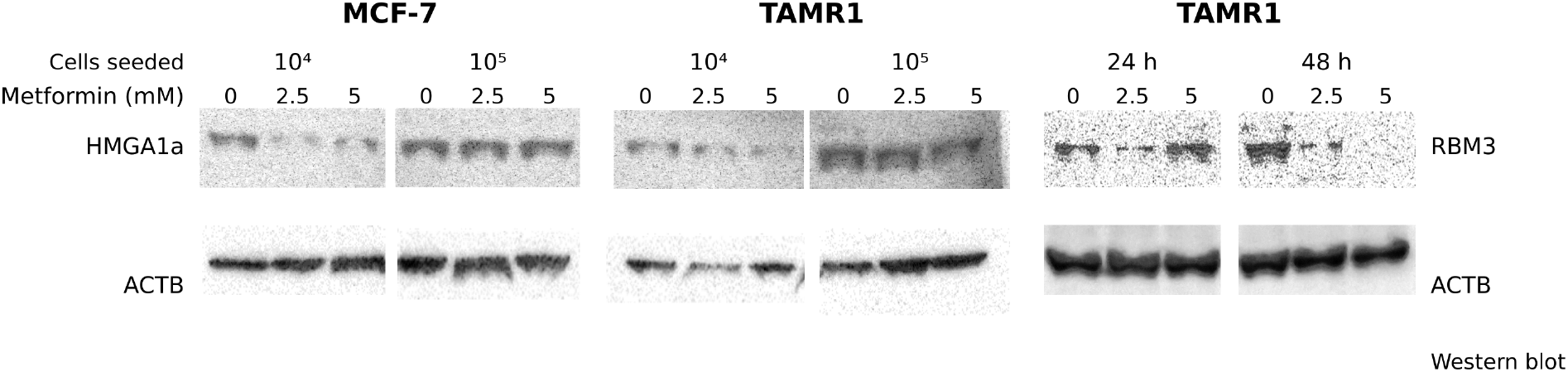
Metformin reduces HMGA1a and RBM3 protein. Immunoblots of MCF-7 and TAMR1 cell lysates after 24 h of treatment with 0, 2.5 or 5 mM metformin at two seeding densities (10^4^ and 10^5^ cells), probed for HMGA1a, and of TAMR1 lysates at 24 and 48 h probed for RBM3, with ACTB as loading control. The four HMGA1a panels are regions of one membrane and the ACTB panels come from four separate images; uncropped membranes are provided as Supplementary Figure S1. Single experiment.

The direction of the HMGA1a result is the opposite of what the splicing data would predict. HMGA1a generates ERα46 by trapping U1 snRNP at an upstream pseudo-5′ splice site [8,9], so less HMGA1a should mean less ERα46 and a higher full-length share. In TAMR1 cells at low density, metformin lowers the full-length share [11] while also lowering HMGA1a.

### ESR1 transcripts in the xenograft tumours

RNA extracted from the xenograft tumours of the earlier study [11] was analysed by RT-PCR for full-length *ESR1* , ERα46 and ACTB (Fig. 4). In MCF-7 tumours, both products were detected in the vehicle group and the ERα46 product alone in the metformin group, while neither product was detected in the tamoxifen or combination groups. In TAMR1 tumours, the full-length product was detected in the vehicle and combination groups but not in the tamoxifen or metformin groups, and the ERα46 product was detected in all four groups.

**Figure 4.**
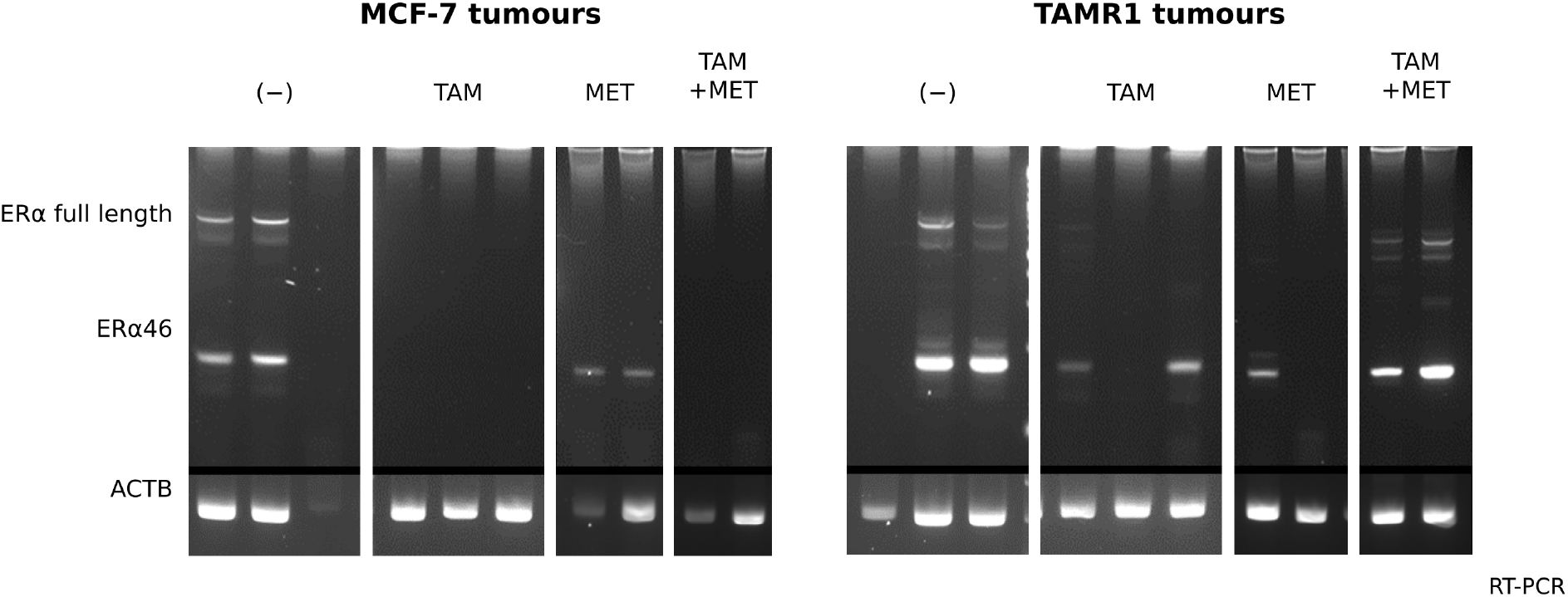
*ESR1* transcripts in the xenograft tumours. RT-PCR for full-length *ESR1* , ERα46 and ACTB on RNA extracted from the MCF-7 (left) and TAMR1 (right) xenograft tumours of the earlier study [11], grouped by treatment: vehicle (−), tamoxifen (TAM), metformin (MET) or both. The *ESR1* and ACTB panels of each group are separate gels and are separated by a horizontal rule. Within each treatment group, each lane is one tumour. The mice carried an MCF-7 tumour and a TAMR1 tumour on opposite flanks and were identified by ear mark, but the source records do not state which animal and which flank each lane came from, so the lanes cannot be matched to individual tumours or paired between the two panels. Lanes in which amplification failed are shown as obtained.

Amplification failed in a number of lanes, including lanes in which ACTB amplified, so these panels cannot be read quantitatively. The full-length product was present in the untreated TAMR1 tumours as well as in the combination group, so these data do not show that the drug combination specifically induces full-length ERα.

## Discussion

The result that matters most here is a negative one. The opposite-direction shift in *ESR1* splicing that we reported at 2.5 and 5 mM [11] does not appear when the same two concentrations are tested at high seeding density, and the four lower concentrations produce no graded effect in either line. A pharmacological effect on splicing would normally show some relation to dose; this one does not. The published observation should therefore be read as holding for the culture condition in which it was made, not as a general property of metformin in these cells.

Cell density is a plausible reason for the discrepancy rather than an incidental one. ERα46 expression is known to rise when MCF-7 cells become hyperconfluent, a limitation we noted in the earlier report [11], so the baseline ratio against which metformin is measured is itself density-dependent. The present data show that the direction of the metformin effect, and not only its size, changes with density.

There is also a precedent for a splicing factor whose nuclear abundance is set by cell density. In MCF10A mammary epithelial cells, nuclear PTBP1 falls at high density and rises at low density, mirroring the behaviour of YAP, and the same holds for cell area and matrix stiffness; blocking proliferation does not change it, so the effect follows the mechanical environment rather than growth rate [12]. Mechanical control of splicing has since been described for several other RNA-binding proteins, among them hnRNPC and RBM4 [13,14]. A splicing outcome that reverses with seeding density is therefore not anomalous, and the density at which a splicing measurement is made belongs in the methods of any such experiment. We did not measure PTBP1 here, and we do not claim it is involved in *ESR1* exon 1 skipping; the point is that the class of mechanism exists.

We asked whether the same holds at the mRNA level by reanalysing the public confluency RNA-seq of Lobo et al. [16] (Supplementary Figure S2). The density contrast behaves as expected: the YAP target genes CTGF and CYR61 fall at high density in all three cell lines (log_2_ fold change −3.21, −2.14 and −0.68, and −1.90, −1.84 and −1.47), as does MKI67. Against that background, PTBP1 mRNA is lower at high density in all three lines (−0.30, −1.49 and −0.07; combined p = 3.7 × 10^−10^), which agrees in direction with the protein and localisation data of Tseng et al. [12]. The effect size differs more than twentyfold between lines, so we would not describe it as uniform.

RBM3 and HMGA1 do not behave this way. RBM3 falls at high density in A375 (−0.59) but not in HEK293 (+0.06) or SH-SY5Y (−0.06), and HMGA1 falls in A375 (−0.87) and SH-SY5Y (−0.19) while rising in HEK293 (+0.20). Neither is consistent across the three lines. We report this because our first pass through the same data gave the opposite impression: using the deposited matrix as supplied, RBM3 appeared to fall in all three lines with a combined p of 0.039. That matrix is not centred between samples, and the apparent effect was an artefact of the scaling rather than a property of RBM3; it disappeared on renormalisation. We mention it as a caution about reusing deposited count matrices without checking them, and because the uncorrected result would have supported our own argument.

These cells are not breast epithelium and none of the three expresses *ESR1* at a measurable level (mean scaled counts 0.1 to 5.4), so this reanalysis bears only on the abundance of the candidate factors, not on *ESR1* splicing.

We also looked for public data that would test the central question directly, and there are none. All 80 human metformin RNA-seq series in the Gene Expression Omnibus were examined: none uses an oestrogen receptor-positive breast cancer line. The two breast datasets are MDA-MB-231, which is triple negative and does not express *ESR1* . MCF-7 cells treated with 1 mM metformin do appear in a 34-compound screen [17], but on a 3′-biased gene-level array, and the difference between full-length ERα and ERα46 lies at the 5′ end of the transcript, so that platform cannot resolve it in principle. Even with suitable raw data, short-read transcript quantification separates 5′-variant isoforms poorly, and *ESR1* carries several promoter-derived 5′ variants. The nested RT-PCR used here, with a primer inside the upstream E/F exon, is the more direct measurement. Whether metformin alters *ESR1* splicing is therefore a question that currently requires new sequencing, for which long reads would be better suited than short ones.

The HMGA1a result argues against the mechanism we had assumed. HMGA1a generates ERα46 by trapping U1 snRNP at an upstream pseudo-5′ splice site [8,9], so lower HMGA1a predicts less ERα46. Metformin lowered HMGA1a in both lines, yet in TAMR1 cells at low density it raises ERα46 [11]. HMGA1a abundance cannot account for that, and we do not propose it as the mediator.

The fall in HMGA1a itself is not an isolated observation. Metformin has been reported to lower nuclear HMGA1 in a germ cell tumour line within 12 h, dose-dependently up to 5 mM, with downstream effects on cyclin D1, the IGF system and MMP-11 [18] — the same direction and the same concentration range as here, in an unrelated cell type. Whether this depends on AMPK was not tested in that study, and we did not test it either. A route that does not pass through AMPK is equally available: HMGA1 is an established nuclear target of insulin receptor signalling, phosphorylated on its C-terminal tail in response to insulin with consequent loss of DNA binding [19]. We make no claim about which route operates here.

RBM3 is the better candidate on present evidence. Metformin lowers RBM3 and alters splicing through it in myotonic dystrophy [10], and we find RBM3 falls with metformin in TAMR1 cells. This remains a correlation: the test is knockdown of RBM3 in TAMR1 cells with the full-length/ERα46 ratio as readout, at a stated seeding density, which we have not done.

The direction of the RBM3 change also needs comment. Nuclear RBM3 is an independent favourable prognostic factor in breast cancer, associated with longer overall and recurrence-free survival, with low grade, small tumour size, ER positivity and Ki-67 negativity, and the association holds after controlling for tamoxifen treatment [15]. Metformin lowered RBM3 in TAMR1 cells here, which is the opposite of the direction that marker would predict for a favourable outcome, while metformin reduced TAMR1 tumour weight in the same model [11]. We cannot reconcile the two from the present data. RBM3 abundance in a cultured cell line and nuclear RBM3 staining in a patient tumour are not the same measurement, and a prognostic association is not a statement about what lowering RBM3 does. The experiment that would settle whether RBM3 carries the splicing effect is the knockdown described above.

The tumour RT-PCR does not resolve why metformin reduced TAMR1 tumour weight while the combination with tamoxifen did not [11]. The full-length product appeared in the untreated tumours as well as in the combination group and failed to amplify in the single-agent groups, so the panels cannot separate the treatments. A quantitative assay against a reference gene, on RNA of confirmed quality, would be needed to ask the question properly.

### Limitations

The dose-response gel is a single experiment and its band ratios were re-measured from the gel image for this report, because the original quantification could not be located. The 48 h panels are single experiments, and the TAMR1 48 h gel yielded one interpretable lane. The immunoblots are single experiments without densitometry. The tumour RT-PCR is qualitative and failed in several lanes. Only one tamoxifen-resistant subline was examined. A negative dose-response from one gel does not overturn a published positive result; it shows that the result does not hold under the conditions tested here and that the effect needs to be re-established with the seeding density controlled and stated.

The public-data reanalysis carries its own limits. It rests on one deposited dataset with three replicates per condition, in three cell lines that are not breast epithelium and do not express *ESR1* , and it measures mRNA where the published density effect on PTBP1 was shown for protein and subcellular localisation. It therefore supports the existence of density-dependent regulation of a splicing factor and says nothing about *ESR1* . The search for metformin datasets was confined to the Gene Expression Omnibus and to series indexed as human RNA-seq; a relevant dataset deposited elsewhere, or indexed differently, would have been missed.

## Supporting information

Supplementary Figure S1. Uncropped gels and blots

Supplementary Figure S2. Confluency effect on candidate factors (GSE249290)

Source Data for Figure 1. Densitometry values

Source Data for Supplementary Figure S2. Candidate factor summary

Source Data for Supplementary Figure S2. All tested genes

Analysis code for the reanalysis of GSE249290

## Declarations

### Ethics approval

All animal protocols followed National Institutes of Health guidelines and were approved by the Fukuoka University Committee for Animal Care (Protocol No. 171116, accepted April 2019).

### Relation to previous work

The cells, compounds and animals used here are those of Harada et al., Am J Pharm Health Res 2022;10(11) [11]. The data shown in Figures 1 to 4 of the present report do not appear in that paper. Conversely, the *ESR1* RT-PCR at 24 h and low density, the ERα immunoblots with and without tamoxifen, the cell viability assays and the tumour and body weights are published there and are not reproduced here.

### Author contributions

Conceptualization: K.O. Methodology: K.O. Investigation: Y.Ha. performed the cell culture, RT-PCR and immunoblot experiments; T.T. and Y.Hm. performed the mouse experiments; K.O. Formal analysis: K.O. Visualization: K.O. Writing — original draft: K.O. Supervision and project administration: K.O. Y.Ho., H.Ha., M.M. and Y.M. advised on study design and interpretation. Writing — review and editing: all authors. All authors read and approved the final manuscript. (Initials: Y.Ha., Yoshihiro Harada; T.T., Tomoko Tanaka; Y.Hm., Yuriko Hamaguchi; Y.Ho., Yuta Horita; H.Ha., Hiroyoshi Harada; M.M., Masayoshi Mori; Y.M., Yusuke Murata; K.O., Kenji Ohe.)

### Data availability

Densitometry values for Figure 1 are provided as a source-data file. Uncropped images of the blots and gels are provided as Supplementary Figure S1. The reanalysis in Supplementary Figure S2 uses GSE249290 [16], which is publicly available from the Gene Expression Omnibus; the renormalised per-gene output and the analysis code are provided with this report.

---

**Supplementary Figure S2. Confluency effect on candidate factors in public RNA-seq (GSE249290)**. Log_2_ fold change, high density versus low density, for HEK293, A375 and SH-SY5Y cells, with 95% confidence intervals (n = 3 per condition). Counts were renormalised by median-of-ratios before testing; the deposited matrix is not centred between samples. The lower block holds the YAP target genes CTGF and CYR61, the Hippo effector YAP1 and the proliferation marker MKI67, which serve as controls that the density contrast is present. Positive values indicate higher expression at high density.

