## Supplementary Figure S1. Uncropped gels and blots for "Metformin lowers HMGA1a and RBM3 protein in breast cancer cells, and its effect on *ESR1* splicing depends on cell density and exposure time"

### Supplementary Figure S1
### Uncropped gels and blots

Harada et al. Metformin lowers HMGA1a and RBM3 protein in breast cancer cells, and its effect on ESR1 splicing depends on cell density and exposure time

**Figure 1 — ESR1 RT-PCR, metformin dose response (high density), uncropped gel**

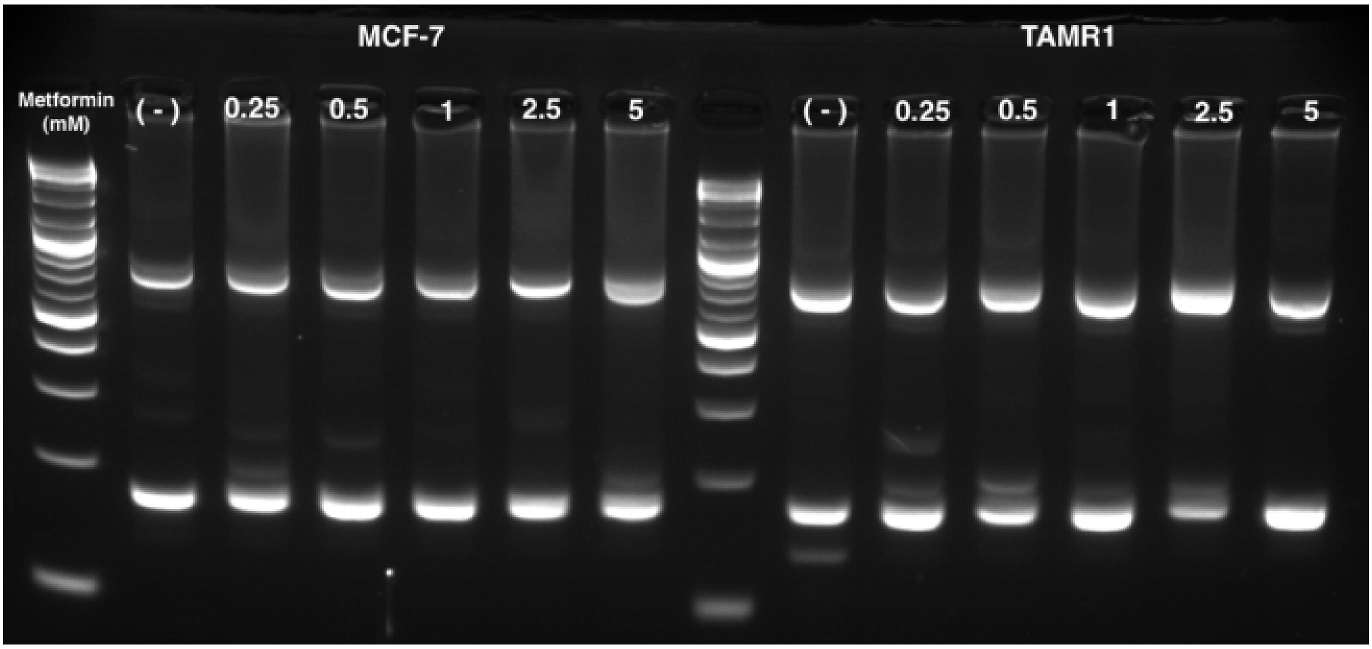


12 sample lanes plus two DNA size markers. Lanes 1-6 MCF-7, lanes 7-12 TAMR1, each 0, 0.25, 0.5, 1, 2.5 and 5 mM metformin.

#### Figure 3 — HMGA1a immunoblot, uncropped membrane

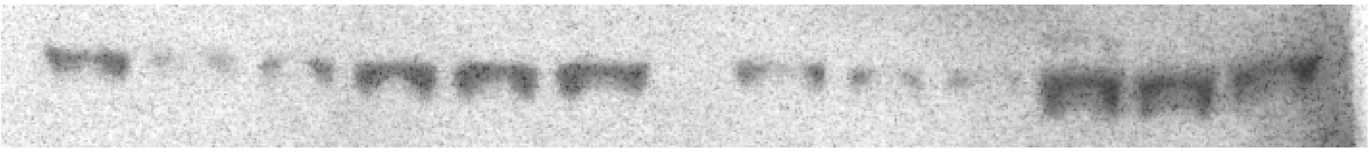

One membrane of twelve lanes. Lanes 1-3 MCF-7 $10^4$ cells, 4-6 MCF-7 $10^5$ cells, 7-9 TAMR1 $10^4$ cells, 10-12 TAMR1 $10^5$ cells, each 0, 2.5 and 5 mM metformin. Figure 3 shows these four groups side by side.

**Figure 3 — RBM3 immunoblot, uncropped membrane**

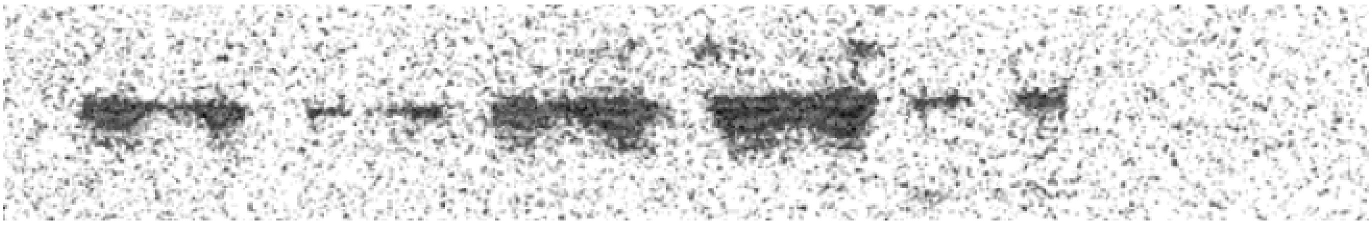

TAMR1 cells, 24 h and 48 h, 0, 2.5 and 5 mM metformin.

**Figure 3 — RBM3 membrane, original scan with size marker**

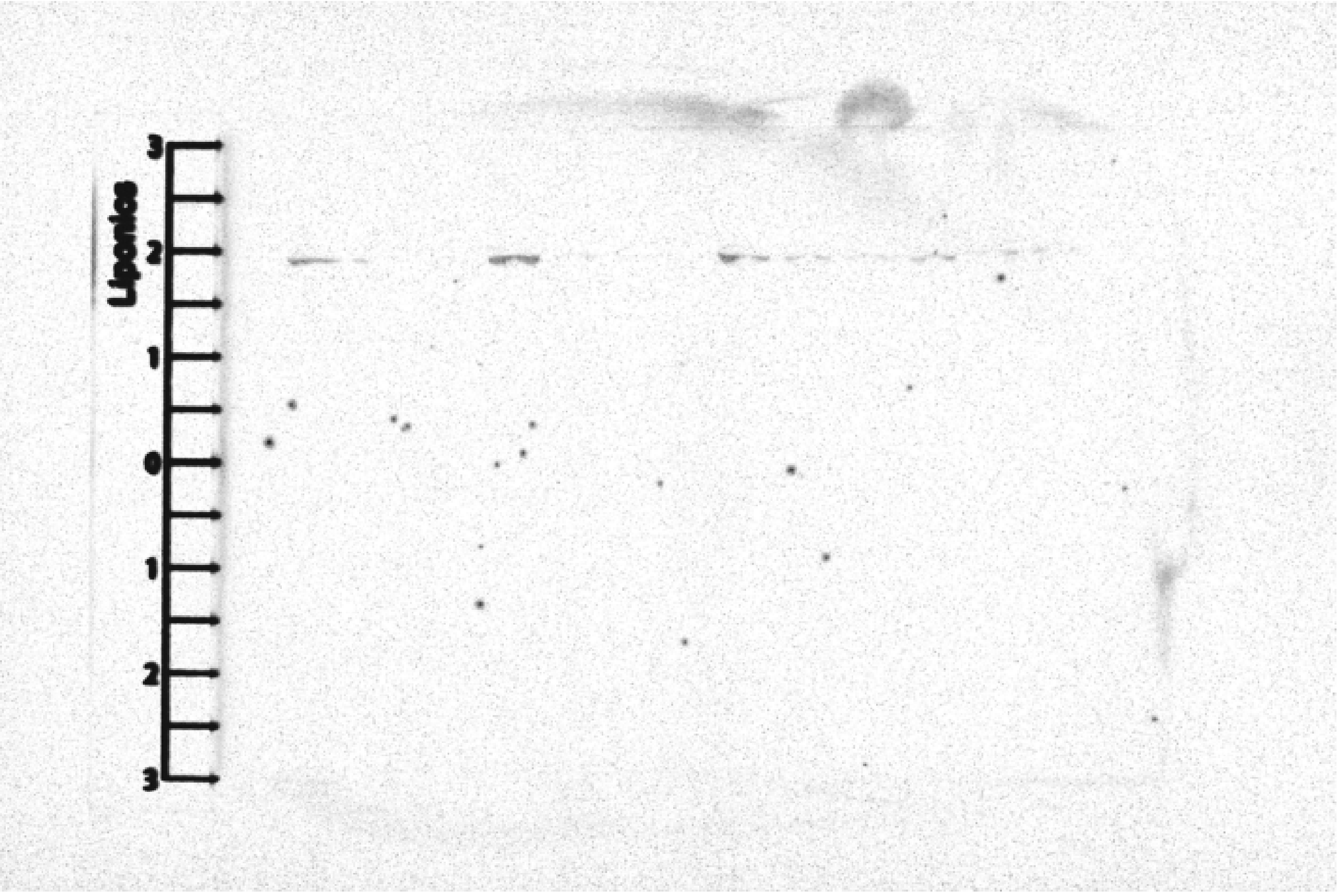


Original scan of the membrane before cropping.

**Figure 2 — ESR1 RT-PCR at low density, the gel regions as supplied**

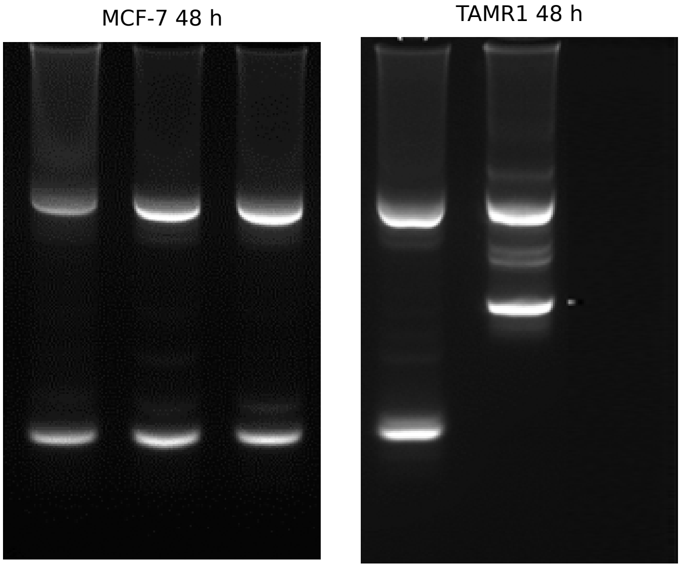


These are the gel regions as they were stored in the source presentation; no further cropping was applied in Figure 2. Only the 0 mM lane of the TAMR1 region gave an interpretable ratio. The corresponding 24 h regions belong to the published experiment and are not shown.

#### **Figure 3 — ACTB loading controls, uncropped images**

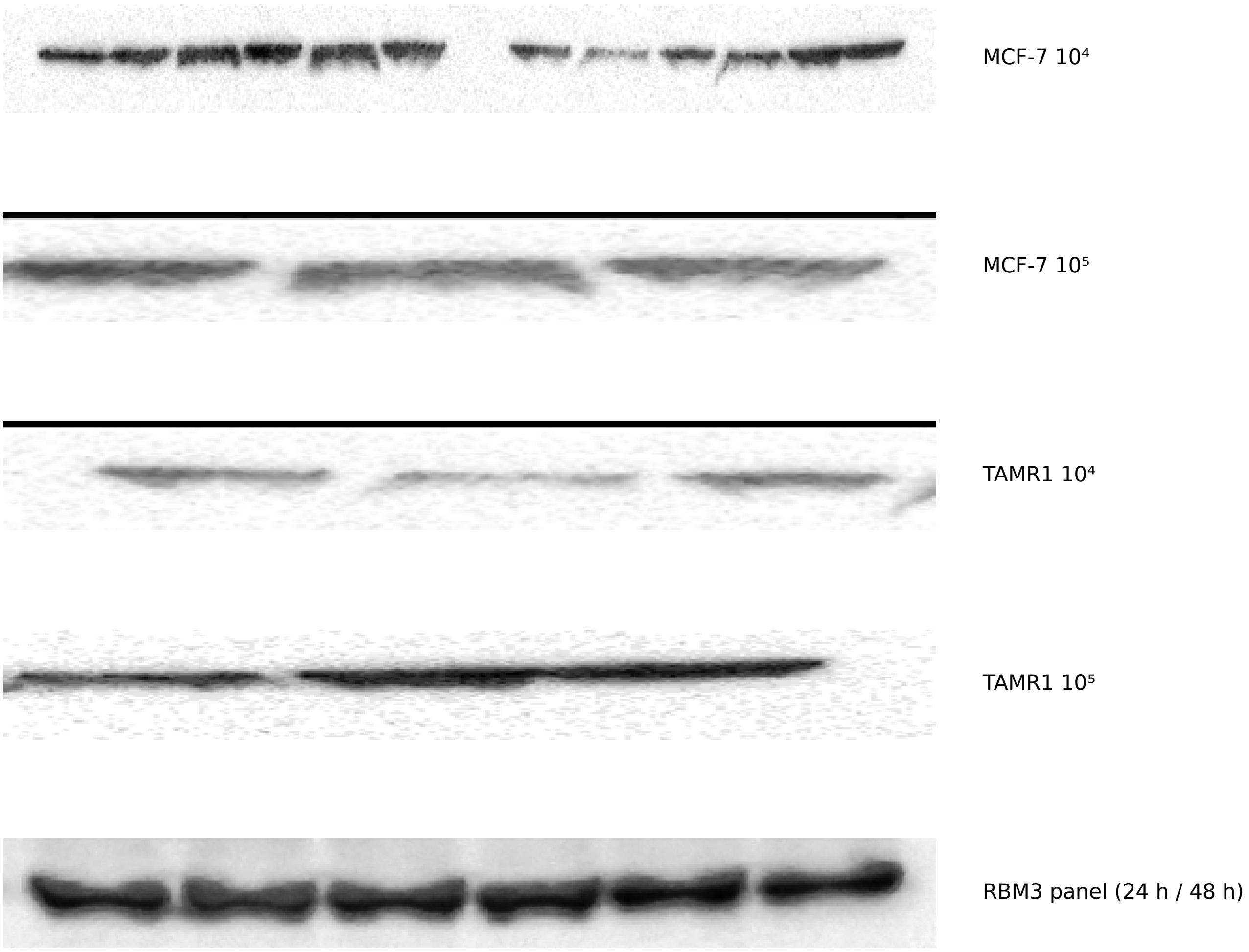


The ACTB panels of Figure 4 come from four separate images rather than from one membrane; each is shown here in full.
