## Supplementary Figure S2. Confluency effect on candidate factors (GSE249290) for "Metformin lowers HMGA1a and RBM3 protein in breast cancer cells, and its effect on *ESR1* splicing depends on cell density and exposure time"

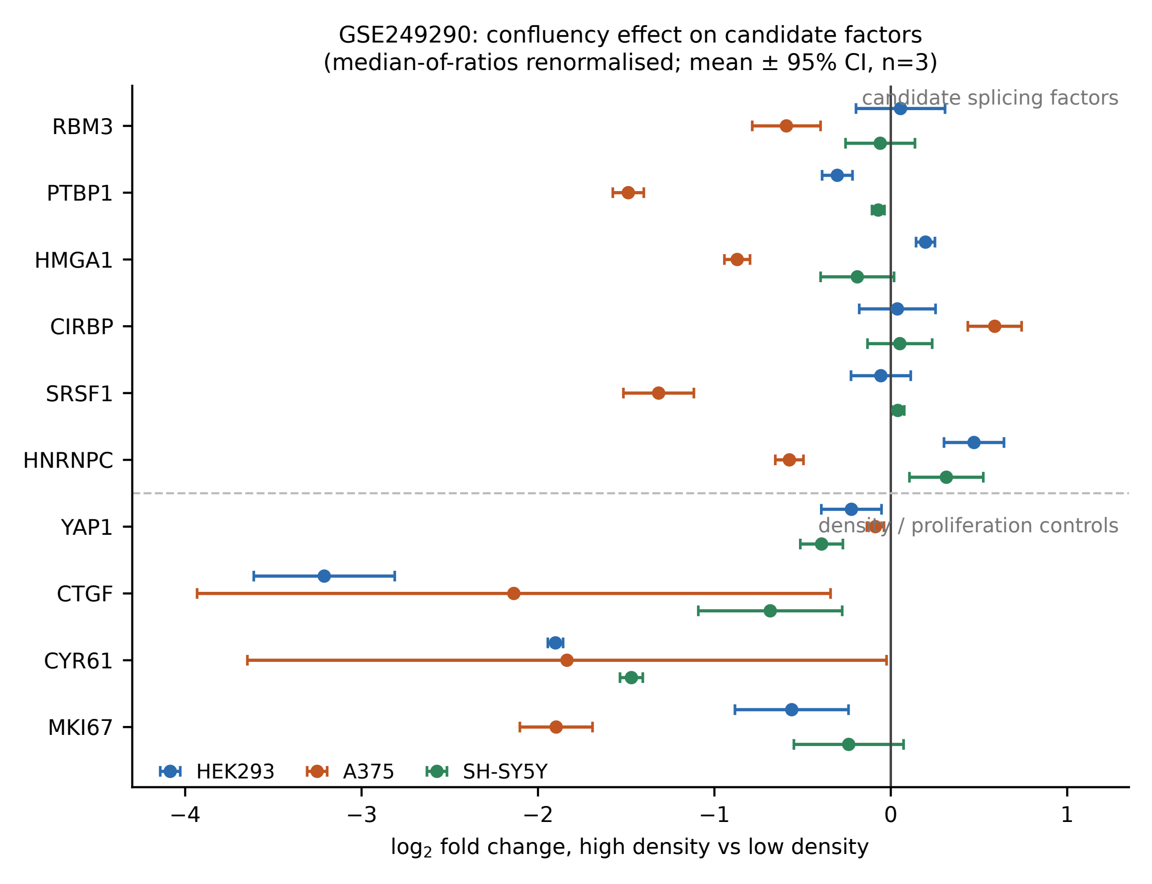

GSE249290: confluency effect on candidate factors
(median-of-ratios renormalised; mean ± 95% CI, n=3)
candidate splicing factors
density / proliferation controls
RBM3
PTBP1
HMGA1
CIRBP
SRSF1
HNRNPC
YAP1
CTGF
CYR61
MKI67
HEK293
A375
SH-SY5Y
−4
−3
−2
−1
0
1
log2 fold change, high density vs low density
